# Thermal acclimation alters the roles of Na^+^/K^+^-ATPase activity in a tissue-specific manner in *Drosophila melanogaster*

**DOI:** 10.1101/2020.11.14.383091

**Authors:** Alexandra Cheslock, Mads Kuhlmann Andersen, Heath A. MacMillan

## Abstract

Insects, like the model species *Drosophila melanogaster*, lose neuromuscular function and enter a state of paralysis (chill coma) at a population- and species-specific low temperature threshold that is decreased by cold acclimation. Entry into this coma is related to a spreading depolarization in the central nervous system, while recovery involves restoration of electrochemical gradients across muscle cell membranes. The Na^+^/K^+^-ATPase helps maintain ion balance and membrane potential in both the brain and hemolymph (surrounding muscles), and changes in thermal tolerance traits have therefore been hypothesized to be closely linked to variation in the expression and/or activity of this pump in multiple tissues. Here, we tested this hypothesis by measuring activity and thermal sensitivity of the Na^+^/K^+^-ATPase at the tagma-specific level (head, thorax and abdomen) in warm-(25°C) and cold-acclimated (15°C) flies by Na^+^/K^+^-ATPase activity at 15, 20, and 25°C. We relate differences in pump activity to differences in chill coma temperature, spreading depolarization temperature, and thermal dependence of muscle cell polarization. Differences in pump activity and thermal sensitivity induced by cold acclimation varied in a tissue-specific manner: While cold-acclimated flies had decreased thermal sensitivity of Na^+^/K^+^-ATPase that maintains activity at low temperatures in the thorax (mainly muscle), activity instead decreased in the heads (mainly brain). We argue that these changes may assist in maintenance of K^+^ homeostasis and membrane potential across muscle membranes and discuss how reduced Na^+^/K^+^-ATPase activity in the brain may counterintuitively help insects delay coma onset in the cold.

## Introduction

Chill susceptible species suffer from cold-induced physiological disruptions that cause tissue damage and death well above the freezing point of their bodily fluids (Bale, 1996; Overgaard and MacMillan, 2017). Thermal limits of these insects can be measured by quantifying chilling injuries or failure to maintain physiological homeostasis in the cold. One such measure is the temperature leading to the progressive onset of chill coma. Neuromuscular failure in the cold is initiated by the insect first losing the ability to perform coordinated movements (at a temperature called the critical thermal minimum; CT_min_), which is immediately followed by complete neuromuscular paralysis (chill coma onset; CCO) (Hazell and Bale, 2011; Mellanby, 1939; Overgaard and MacMillan, 2017; Sinclair et al., 2015). This paralytic state is reversible, and when returned to permissive temperatures the insect will recover (quantified as the chill coma recovery time) (David et al., 1998; Overgaard and MacMillan, 2017).

The onset of cold-induced coma has been causally linked to a shutdown of central nervous system (CNS) function (Robertson et al., 2017). The CNS is separated from the hemolymph by the blood brain barrier which regulates ion entry via selective ion channels and transporters, resulting in maintenance of a relatively low local extracellular [K^+^] within the interstitium (≤ 10 mM) (Andersen et al., 2018; Armstrong et al., 2012; Limmer et al., 2014). Cold exposure, however, tends to slow and ultimately disrupt ionoregulatory mechanisms (Koštál et al., 2006; MacMillan and Sinclair, 2011a; Overgaard and MacMillan, 2017). In the insect CNS this leads to a phenomenon known as spreading depolarization (SD), which is characterized by a rapid surge in [K^+^] in the extracellular space surrounding neural and glial cells (30 - 60 mM, depending on species), followed by complete silencing of the CNS (Andersen et al., 2018; Armstrong et al., 2012; Robertson et al., 2020, 2017).

While a loss of ion balance occurs rapidly within the CNS and leads to the coma phenotype, cold-induced mortality is hypothesized to be caused by gradual, loss of systemic (hemolymph) K^+^ balance (Koštál et al., 2006, 2004; MacMillan and Sinclair, 2011b; Overgaard and MacMillan, 2017). At permissive temperatures, hemolymph ion balance is regulated by balancing passive diffusion with active transport in the renal system (Beyenbach et al., 2010; Phillips et al., 1986). In the cold, however, renal function is compromised and ions will tend to move down their respective electrochemical gradients (Andersen and Overgaard, 2020; Des Marteaux et al., 2018; Gerber and Overgaard, 2018; Koštál et al., 2006; MacMillan and Sinclair, 2011b; Yerushalmi et al., 2018). Consequently, hemolymph [Na^+^] and volume tend to decrease, while hemolymph [K^+^] gradually increases (hyperkalemia), resulting in depolarization of cells bathed in the hemolymph (e.g. muscle; Overgaard and MacMillan, 2017). This systemic cold- and hyperkalemia-induced depolarization is thought to initiate cell death, particularly in the muscle tissue and ultimately leading to organismal chilling injury (Bayley et al., 2018; Carrington et al., 2020; MacMillan et al., 2015b). If an insect is removed from the cold before extensive injury occurs, successful recovery from chill coma is dependent on the rate of hemolymph ion and water balance restoration (MacMillan et al., 2012), which is tightly linked to the capacity of the renal system to restore hemolymph volume, and Na^+^ and K^+^ balance (MacMillan et al., 2014; Overgaard and MacMillan, 2017).

Notably, chill tolerance phenotypes are highly plastic in many insects, including *D. melanogaster*, and rapid cold-hardening (on the order of minutes to hours) or cold acclimation (on the order of days to weeks) can allow chill susceptible insects to mitigate the effects of chilling on the brain (Armstrong et al., 2012), maintain renal function at low temperatures (Andersen and Overgaard, 2020; Yerushalmi et al., 2018), prevent hemolymph hyperkalemia (MacMillan et al., 2015a), avoid Ca^2+^ overload and muscle cell death (Andersen et al., 2017a; Bayley et al., 2020; MacMillan et al., 2017), or more rapidly restore ion balance upon rewarming (Findsen et al., 2013). If insects can prevent some or all of these effects of chilling through phenotypic plasticity, organismal chill tolerance improves, and these protective effects are often directly associated with modulation of ion homeostasis.

While there is ample evidence to suggest a direct role of ionoregulatory physiology in modulating chill susceptibility, the role of specific ionoregulatory mechanisms in each organ (and therefore each cold tolerance phenotype) remains largely unknown. However, the key role of both local (CNS) and systemic (hemolymph) hyperkalemia in limiting organismal function (i.e. chill coma and injury phenotypes), and the ability of cold-acclimated insects to mitigate these effects suggests a central role of Na^+^/K^+^-ATPase activity in chill susceptibility (MacMillan and Sinclair, 2011a). This ATP-dependent pump greatly contributes to the maintenance of ion gradients within and across most animal tissues, and its impaired function has been implicated in the loss of ion balance caused by multiple environmental stressors (Boutilier, 2001; Rodgers et al., 2010; Spong et al., 2016a). Despite an apparent role for the pump in stress tolerance, direct evidence that modifications to Na^+^/K^+^-ATPase activity underlie variation in chill tolerance traits is scarce and what evidence exists is confusing. At the whole animal level, MacMillan et al. (MacMillan et al., 2015c) found reduced Na^+^/K^+^-ATPase activity and no change in thermal sensitivity of the pump in cold-acclimated flies. Similarly, Yerushalmi et al. (2018) found lower Na^+^/K^+^-ATPase activity in the gut and Malpighian tubules of cold-acclimated *D. melanogaster* despite these flies having higher rates of ion transport across the Malpighian tubule and rectal epithelia that their warm-acclimated conspecifics (Yerushalmi et al., 2018). By contrast, Des Marteaux et al. (Des Marteaux et al., 2018) found increased Na^+^/K^+^-ATPase activity, but decreased fluid transport rates in the Malpighian tubules of cold acclimated crickets (*Gryllus pennsilvanicus*). Thus, how insects modulate activity of the Na^+^/K^+^-ATPase in response to cold acclimation remains unresolved, and because the roles of this pump in ionoregulation vary according to tissue, different tissues are likely to respond to thermal acclimation in different ways.

Here, we perform a comparative analysis of the differences in Na^+^/K^+^-ATPase activity and thermal sensitivity in cold- and warm-acclimated *D. melanogaster*. Given the diverse roles of individual organs in cold tolerance, and the diverse roles of Na^+^/K^+^-ATPase in cell membrane and epithelial transport processes within these organs, we hypothesized that adaptive changes in pump function induced by cold acclimation would be organ-specific. To test this, we measured maximal activity of Na^+^/K^+^-ATPase in each of the three tagmata (head, thorax, abdomen) at three temperatures (15, 20 and 25°C), in order to detect any possible alteration in thermal sensitivity between cold- and warm-acclimated flies. We specifically hypothesized that Na^+^/K^+^-ATPase activity in the thorax (mainly muscle) of cold-acclimated flies would be better maintained at low temperature as this would help maintain muscle membrane potential (i.e. excitability). Lastly, we hypothesized that the head (mainly CNS) of cold acclimated flies would have increased Na^+^/K^+^-ATPase activity and/or a decrease in thermal sensitivity of pump activity, either of which should help prevent cold-induced SD (i.e. onset of chill coma) if the Na^+^/K^+^-ATPase is directly involved in this phenomenon.

## Materials and Methods

### Animal husbandry

The *Drosophila melanogaster* used in our experiments originated from isofemale lines collected in London, and Niagara on the Lake, ON, Canada (Marshall and Sinclair, 2010). All flies were reared on a banana-based diet at a population density of approximately 150 – 200 flies per 180 mL bottle. Flies were raised from egg to adult at 25 ± 0.5°C on a 12 h:12 h light:dark cycle in an incubator (Precision 818; Thermo Fisher Scientific, Ottawa, ON, Canada) with humidity maintained between 40 and 60%. To raise experimental adults, eggs were collected from adult flies for a period of 2.5 – 3 h to achieve a density of ~150 eggs per bottle before the adults were removed. Newly eclosed flies were transferred to a separate bottle within 24 h to control their age and held at 25°C. Three days after this separation, the flies were lightly anesthetized with a brief (approx. 5 min) carbon dioxide exposure and sexed. Males were discarded and females were collected, separated into smaller groups of approximately 50 flies per 40 mL vial for acclimation. Half of these vials were returned to the 25°C incubator (warm acclimation) while the rest were placed into a separate incubator (MIR-154-PA incubator; Panasonic Corporation, Kadoma, Japan) for cold acclimation at 15°C with an identical light:dark cycle. Female flies were used for experiments after seven days of acclimation (total age of 10 ± 1 days).

### Chill coma onset

To quantify the chill coma onset temperature, female flies from each acclimation group were individually aspirated into 3.7 mL glass screw-top vials, and subsequently randomized to ensure that the experimenter was blinded with regards to acclimation treatment. All vials were affixed to a custom-built aluminum rack and submerged into an aquarium (61.0 × 21.1 × 51.7 cm) filled with a 1:1 mixture of water and an ethylene glycol-based coolant. This aquarium was connected to a refrigerated circulator (Model AP28R-30, VWR International, United States) which circulated the coolant at a pre-set temperature of 20°C. Temperature of the coolant was monitored by averaging the temperature reading of three type-K thermocouples connected to a TC-08 unit and computer running Picolog software (Pico Technologies, Cambridgeshire, UK). Flies were held at 20°C for 15 min after which temperature was decreased at a rate of 0.1°C min^−1^ while being observed. A metal rod was used to systematically tap the vials, and the chill coma onset temperature was quantified as the temperature at which the animal displayed a complete lack of movement in response external stimulus (i.e. tapping the vial). A total of 13 and 14 cold- and warm-acclimated flies were sampled.

### Na^+^/K^+^-ATPase activity

To quickly dissect flies, they were aspirated into a centrifuge tube and snap frozen in liquid nitrogen, after which legs and wings were removed before the flies were quickly separated into tagma (head, thorax, and abdomen). Final samples for homogenization were composed of 25 of each tagma and were stored at −80°C.

Activity of Na^+^/K^+^-ATPase was measured at 15, 20, and 25°C using an enzymatic spectrophotometric assay in samples of tagma collected from both warm- and cold-acclimated flies. The Na^+^/K^+^-ATPase assay we used was a modified 96-well microplate assay described by McCormick & Bern (McCormick and Bern, 1989) and modified by Jonusaite *et al.* (Jonusaite et al., 2011), which relies on measuring the depletion of NADH which is linked to ATP hydrolysis via lactate dehydrogenase and pyruvate kinase. To run the assay, samples of each tagma were first thawed on ice in 800 μL of homogenization buffer (four-parts SEI: 150 mM sucrose, 10 mM EDTA, 50 mM imidazole; pH 7.3, and one-part SEID: SEI plus 0.5% w/v deoxycholic acid) except for heads where only 500 μL was used. Samples were homogenized manually with polypropylene plastic grinding pestles in centrifuge tubes, after which they were sonicated on ice in 4 x 5-7 second intervals with 10-15 second pauses between intervals to avoid overheating and protein denaturation in the samples. Previous studies then centrifuged the samples to use the supernatant, however, we found that approximately half the activity was lost in the thorax samples using this approach. The remaining activity could be found by resuspending the pellet. After comparing the activity from the crude homogenate to the supernatant and pellet samples, we found no differences in total activity or data quality, and therefore used the crude homogenate instead. Each sample was separated into four aliquots and stored at −80°C until use. On the days of the experiment, solution A (4 units/mL lactate dehydrogenase (Sigma, L2500-10KU), 5 units/mL pyruvate kinase (Sigma, P1506-5KU), 2.8 mM phosphoenolpyruvic acid (Sigma-Aldrich, P3637-500MG), 3.5 mM ATP (Bio Basic, AB0020), 0.22 mM NADH (Bio Basic, NBO642), 50 mM imidazole) and solution B (solution A plus 5 mM ouabain (Sigma-Aldrich, O3125-1G)) were prepared. A 3:1 mixture of solution A or B to a salt solution (189 mM NaCl, 10.5 mM MgCl_2_, 42 mM KCl, 50 mM imidazole) was added to activate each solution. Prior to mixing solution B however, an ADP (Bio Basic, AD0016D) standard curve was generated to evaluate the depletion of ADP in solution A (McCormick and Bern, 1989). 10 μL of each ADP standard was loaded into a 96-well microplate in triplicate, followed by 200 μL of solution A. Absorption at 340 nm was measured from each standard and recorded every minute over a 30 minute period in a microplate spectrophotometer (BioTek Epoch; Vermont, United States). Solution A was only used if a negative linear relationship between the final optical density (OD) and the ADP standard concentration was observed to have a slope between −0.009 & −0.014 OD/nM ADP (see supplementary materials for sample standard curve). Once solution A was deemed suitable, 10 μL of crude sample homogenate of each tagma were loaded in sextuplicate, followed first by the addition of solution B and then solution A (three wells were dedicated to each solution per sample). Absorption at 340 nm was then recorded every minute for a duration of 30 minutes using the same microplate spectrophotometer. This was repeated for six biological replicates across three assay temperatures (15°C, 20°C and 25°C). We attempted to also measure activity at 4°C but failed to get reliable measurements in these small quantities of tissue at such a low temperature. Following each Na^+^/K^+^-ATPase assay, the protein content of each sample was quantified using the Bradford assay (B6916, Sigma Aldrich, Saint Louis, MI, USA) with protein standards made from bovine serum albumin (ALB001.25, Bioshop Canada, Burlington, ON, CA). In a 96-well plate, 5 μL of each protein standard and of crude homogenate samples were loaded into individual wells in triplicate, followed by the addition of 250 μL of Bradford reagent to each well. After 10 minutes of incubation, the absorbance of both samples and protein standards were measured using a spectrophotometer at 595 nm. Sample protein concentrations were then calculated by interpolation from the standard curve. Subsequently, Na^+^/K^+^-ATPase activity was calculated as:

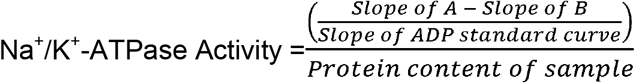

where *Slope of A* refers to the slope produced by activity in solution A (no inhibition), and *Slope of B* refers to the slope produced by activity in solution B (containing ouabain).

For each tagma we calculated the thermal sensitivity (Q_10_) of the Na^+^/K^+^-ATPase as follows:

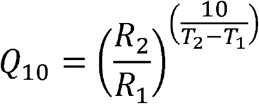

Here, T_2_ and T_1_ are 25 and 15°C, and R_2_ and R_1_ are the mean activities of the Na^+^/K^+^-ATPase at 25 and 15°C, respectively.

### Electrophysiological measurements

To relate Na^+^/K^+^-ATPase activity of the head and thorax to physiological functions we measured the transperineurial potential (TPP; the electric potential generated across the blood-brain-barrier) during cooling from 20°C and muscle membrane potentials (V_m_) at 15-25°C to estimate the thermal sensitivity of the electrogenic effect (which is mainly driven by the Na^+^/K^+^ ATPase, see (Overgaard and MacMillan, 2017)). These measurements were made following a procedure similar to that of Andersen and Overgaard (Andersen and Overgaard, 2019). Briefly, female flies were immobilized in a bed of wax on a glass cover slide (without anesthesia) and placed in a thermoelectrically cooled plate where temperature was monitored with a type-K thermocouple. Here, a small cut was made in the abdomen to insert an Ag/AgCl reference electrode and the fly was prepared for measurement of either TPP or muscle V_m_.

To measure the TPP, a small hole was made in the head using micro scissors and a glass microelectrode was placed into the hemolymph surrounding the brain using a micromanipulator. For these experiments, glass microelectrodes were pulled to a tip resistance of 5-7 MΩ on a Flaming-Brown P-1000 Micropipette puller (Sutter Instruments, Novato, CA, USA). TPP was measured with a Duo 773 intracellular/extracellular amplifier (World Precision Instruments, Sarasota, FL, USA), digitized using a PowerLab 4SP AD converter (ADInstruments Inc., Colorado Springs, CO, USA), and displayed and recorded on a computer. TPP measurements were started at 20°C by zeroing the voltage in the hemolymph around the brain after which the glass electrode was gently inserted into the brain through the blood-brain barrier. This insertion resulted in a small, sudden increase in potential (usually 5-10 mV) and the increase was noted down. Next, temperature was decreased by 1°C min^−1^ while continuously measuring the TPP until a large, abrupt drop in TPP was observed (~ 25-40 mV in ~ 5-10 s), indicative of a spreading depolarization (SD) having occurred. The temperature of the SD was then measured as the temperature of the half-amplitude of the drop in transperineurial potential, and the amplitude was measured as the difference between the transperineurial potential before and after the SD. Only flies where all three parameters (TPP at 20°C, SD temperature, and SD amplitude) were recorded successfully were included in this study, resulting in measurements from six 15°C acclimated flies and seven 25°C acclimated flies.

Muscle membrane potential (V_m_) was measured in the same setup but followed a different approach: After immobilization and insertion of the Ag/AgCl reference electrode, a small hole was made in the top of the thorax with a 26G needle. Next a sharp glass electrode (tip resistance of 10-15 MΩ) was inserted into the flight muscle in the thorax and carefully moved further into the muscle with a micromanipulator. Successful penetration of a muscle cell (and thus measurement of muscle cell V_m_) was indicated by a sharp drop in voltage and when a similar drop in potential could be repeated by retracting and reinserting the electrode three times in quick succession (within 20 s). The muscle cell V_m_ of that cell was then calculated as the average potential drop of the three repeated penetrations. Next, the glass microelectrode was moved to a different muscle cell by changing the angle of approach or by moving further into the thorax to obtain measurements of three different muscle cells. The average fly muscle V_m_ was then calculated as the mean of the three different cells. Muscle V_m_ was measured in seven 15°C acclimated flies and seven 25°C acclimated flies and was measured at both 15 and 25°C in each fly. Measurements started at either 15°C (three flies in each group) or 25°C (four flies in each group), after which the temperature was rapidly changed to the other test temperature (25°C and 15°C, respectively) where muscle V_m_ was measured again after a 10 min waiting period.

### Data analysis

All data analysis was performed in R version 4.0.2 (R Development Core Team, 2019). Differences in CCO temperature were analyzed using a Welch two-sample t-test. The effects of assay temperature and acclimation temperature on Na^+^/K^+^-ATPase activity was analyzed using a generalized linear model and an analysis of deviance. The effect of cold acclimation on the transperineurial potential (at 20°C) and the spreading depolarization temperature and amplitude were all analyzed with Student’s t-tests after using F-tests to confirm that both groups had similar variation. The ability to maintain muscle membrane potential in the cold (between 15 and 25°C) in both acclimation groups was analyzed using a linear mixed effects model (using the lme() function in the ‘nlme’ package for R) which included acclimation temperature and test temperature as fixed factors and the individual fly as a random factor. For these experiments, the order of test temperature was randomized, but the order was found to have no effect of our measurements and was excluded from the analysis. In all analyses, the level for statistical significance was set to 0.05. All values reported and presented are means ± standard error of the mean unless stated otherwise.

## Results

### Chill coma onset temperatures

The chill coma onset temperatures were recorded in 15°C and 25°C acclimated flies to verify that the acclimation protocol elicited plasticity in cold tolerance. As expected, the chill coma temperatures recorded for cold- and warm-acclimated flies differed significantly (t_24.6_ = −13.7, *P* < 0.001) with temperatures being 5.5 ± 0.1°C and 8.0 ± 0.1°C, respectively (Fig. 1).

**Figure 1.**
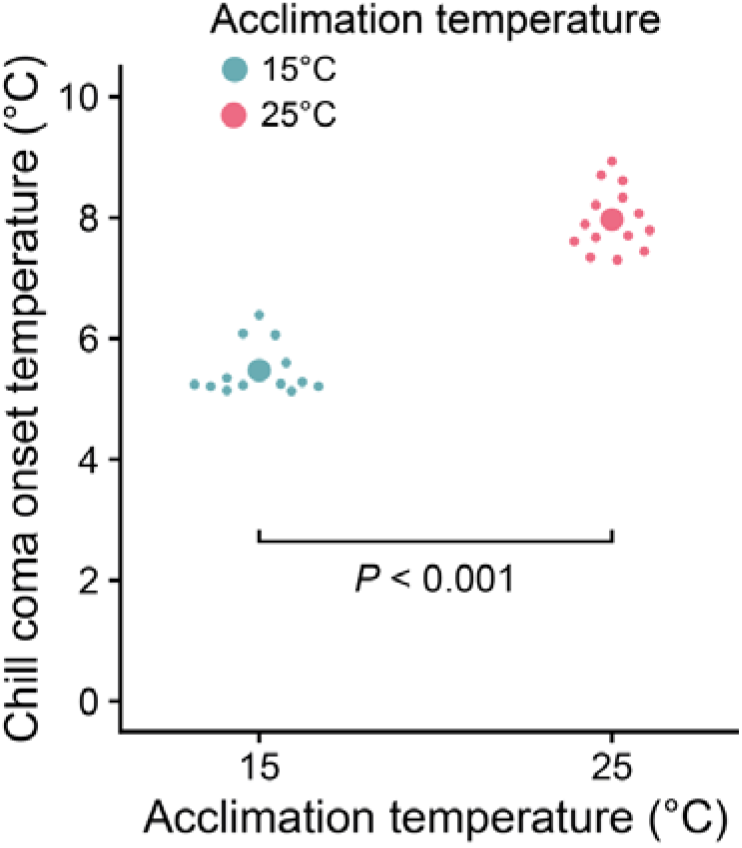
Cold acclimated flies enter chill coma at lower temperatures compared to warm acclimated conspecifics. The chill coma onset (CCO) temperature was measured in 13 cold (15°C, blue) and 14 warm (25°C, red) acclimated *D. melanogaster*. Small circles represent individual observations, and error bars that are not visible are obscured by the symbols.

### Tagma-specific Na^+^/K^+^-ATPase activity and thermal sensitivity

Na^+^/K^+^-ATPase activity was measured in heads, thoraxes, and abdomens of cold-and warm-acclimated flies at 15, 20, and 25°C (**Fig. 2**, see **Fig. S1** for the abdomen data). Na^+^/K^+^-ATPase activity varied between tissue types, with mean activity being the highest in the head samples (e.g. 4.36 μmoles ADP mg protein^−1^ h^−1^, **Fig. 2a**) and lowest in abdomen samples (e.g. 0.32 μmoles ADP mg protein^−1^ h^−1^; see **Fig. S1**) when compared at 25°C. Abdominal Na^+^/K^+^-ATPase measurements were generally low and variable, and are difficult to interpret given the number and varied roles of organs in this tagma. As such we have opted to include these data in a supplementary figure for reference (**Fig. S1**) but do not discuss them further here. Na^+^/K^+^-ATPase activity in the heads (**Fig. 2a**) of cold-acclimated flies had lower Na^+^/K^+^-ATPase activity than those of warm-acclimated flies (F_1,34_ = 3.2, *P* = 0.039). Additionally, Na^+^/K^+^-ATPase activity decreased with temperature in heads samples from both cold- and warm-acclimated flies (F_1,33_ = 22.4, *P* < 0.001). This effect was similar in both acclimation groups (F_1,32_ = 1.1, *P* = 0.209), but despite this, the *Q*_*10*_ coefficient (derived from means) for cold-acclimated flies (1.70) was ~ 21 % lower than that of their warm-acclimated conspecifics (2.15). In the thorax the overall patterns were similar (**Fig. 2b**): there was also no clear effect of acclimation temperature on Na^+^/K^+^-ATPase activity (F_1,34_ = 0.5, *P* = 0.116) despite thoraxes from cold-acclimated flies generally having higher Na^+^/K^+^-ATPase activity below 25°C. The effect of temperature was still present (F_1,33_ = 1.2, *P* = 0.022), but its’ effect was again similar between acclimation groups (F_1,32_ = 0.2, *P* = 0.281) despite *Q*_*10*_ coefficients being ~ 36 % lower for cold-acclimated flies (1.37) compared to the warm-acclimated group (2.14).

**Figure 2.**
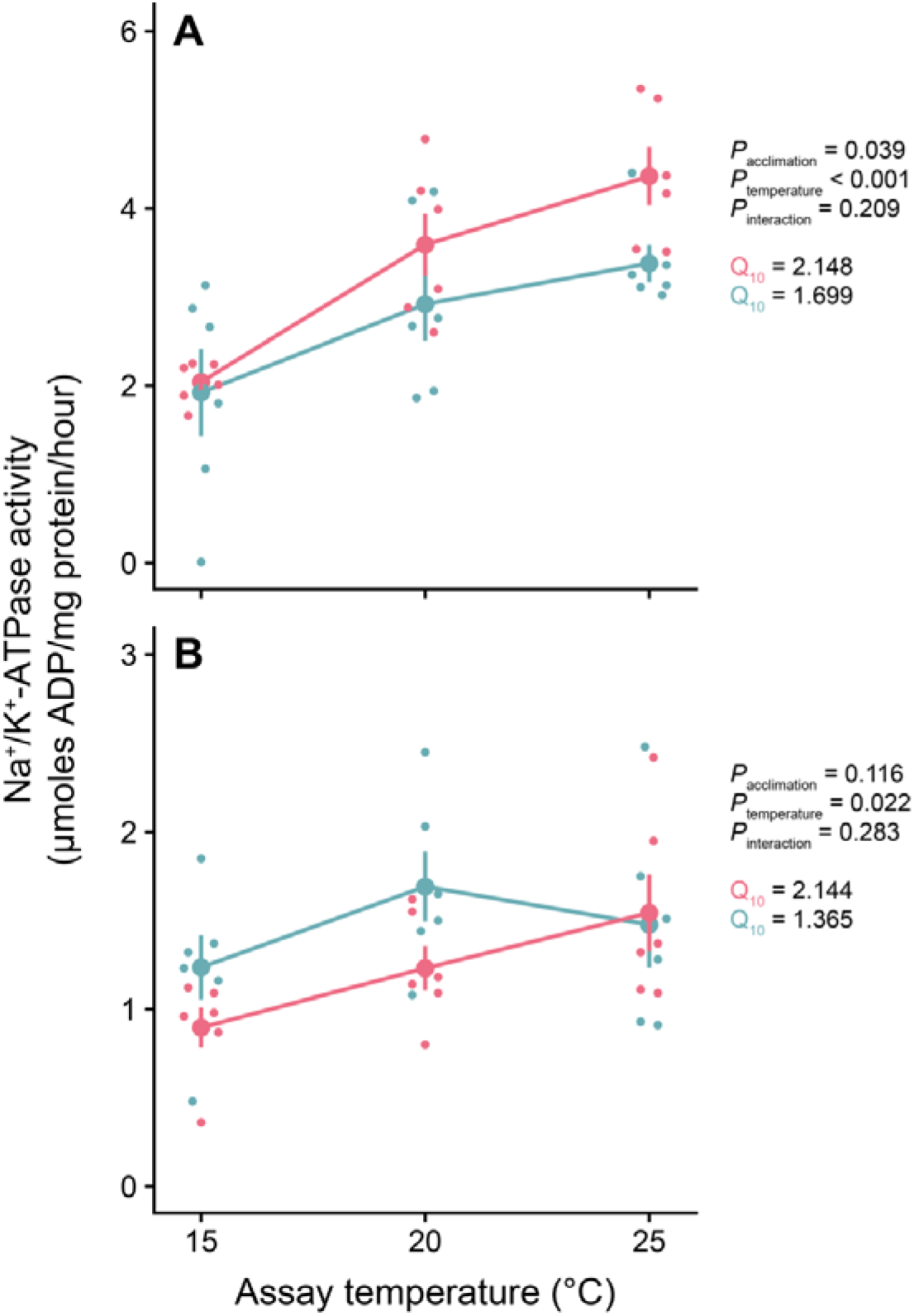
The Na^+^/K^+^-ATPase activity in *D. melanogaster* tissues generally decreases with assay temperature and varies depending on the tagma type and acclimation. Na^+^/K^+^-ATPase activity assays were performed on (A) head, (B) thorax, and abdomen (see Fig. S1) samples collected from 15°C (blue) and 25°C (red) acclimated flies at three temperatures (15, 20 and 25°C). Six biological replicates were sampled per acclimation and tagma combination and were measured at all three assay temperatures. Small circles represent individual observations.

### Transperineurial potential and cold-induced spreading depolarization

The transperineurial potential was recorded at 20°C in both cold- and warm-acclimated flies, after which temperature was decreased by 1°C min^−1^ until a spreading depolarization occurred (**Fig. 3**). Transperineurial potentials differed between acclimation groups when measured at 20°C (t_11_ = 2.5, *P* = 0.031) such that it was higher in the brain of cold-acclimated flies (10.6 ± 1.0 mV) compared to those of warm-acclimated flies (8.1 ± 0.4 mV) (**Fig. 3a**).

**Figure 3.**
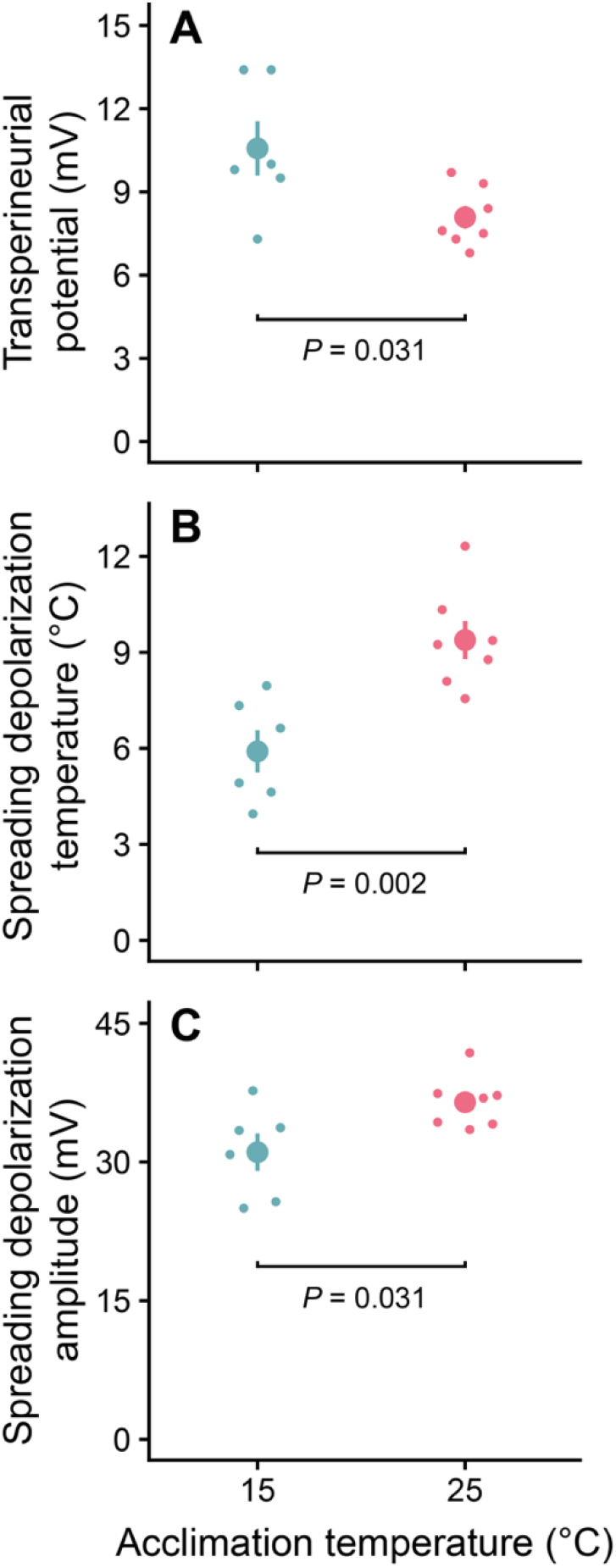
Cold acclimation increases the transperineurial potential and lowers spreading depolarization temperature and amplitude. (A) Transperineurial potential was measured at 20°C in 15°C (blue) and 25°C (red) acclimated *D. melanogaster* after which temperature was lowered by 1°C min^−1^ until a spreading depolarization event was observed, for which the (B) temperature and (C) amplitude were recorded. Small circles represent individual observations, and error bars that are not visible are obscured by the symbols.

The temperature leading to cold-induced SD was recorded as the temperature at the half-amplitude of the SD-induced drop in transperineurial potential. Similar to chill coma onset, cold-acclimated flies had a lower SD temperature than their warm-acclimated conspecifics (t_11_ = −3.9, *P* = 0.002) such that SD happened at 5.9 ± 0.7°C in 15°C acclimated flies and 9.4 ± 0.6 in 25°C acclimated flies (**Fig. 3b**). Interestingly, cold-acclimated flies also had lower SD amplitudes than their warm-acclimated conspecifics (t_11_ = −2.5, *P* = 0.031) such that the drop in TPP in cold- and warm-acclimated flies were 31.1 ± 2.0 mV and 36.5 ± 1.1 mV, respectively (**Fig. 3c**).

### The effect of temperature on muscle membrane potential

Muscle membrane potentials of cold- and warm-acclimated flies were measured in the flight muscle at both 15°C and 25°C in the same individuals (**Fig. 4**). Overall, there was no effect of acclimation (F_1,12_ = 1.4, *P* = 0.262) and muscle V_m_ at 25°C was −64.0 ± 1.1 mV for cold- and −65.2 ± 1.2 mV for warm-acclimated flies. As expected, exposure to 15°C depolarized muscle V_m_ (F_1,12_ = 129.1, *P* < 0.001), but did so in an acclimation-specific manner (F_1,12_ = 12.7, *P* = 0.004) such that muscle V_m_ of cold-acclimated flies depolarized to −57.6 ± 1.2 mV while warm-acclimated flies’ muscle depolarized to −53.0 ± 1.2 mV. The matching measurements at 15°C and 25°C further allowed us to directly quantify the effect of temperature in both acclimation groups (i.e. the interaction between acclimation and assay temperature): The thermal sensitivity differed between the two groups of flies (i.e. the significant interaction above) such that cold-acclimated flies depolarized by 0.6 ± 0.1 mV °C^−1^ while warm-acclimated flies depolarized by 1.2 ± 0.1 mV °C^−1^ when exposed to cold (**Fig. 4, insert**).

**Figure 4.**
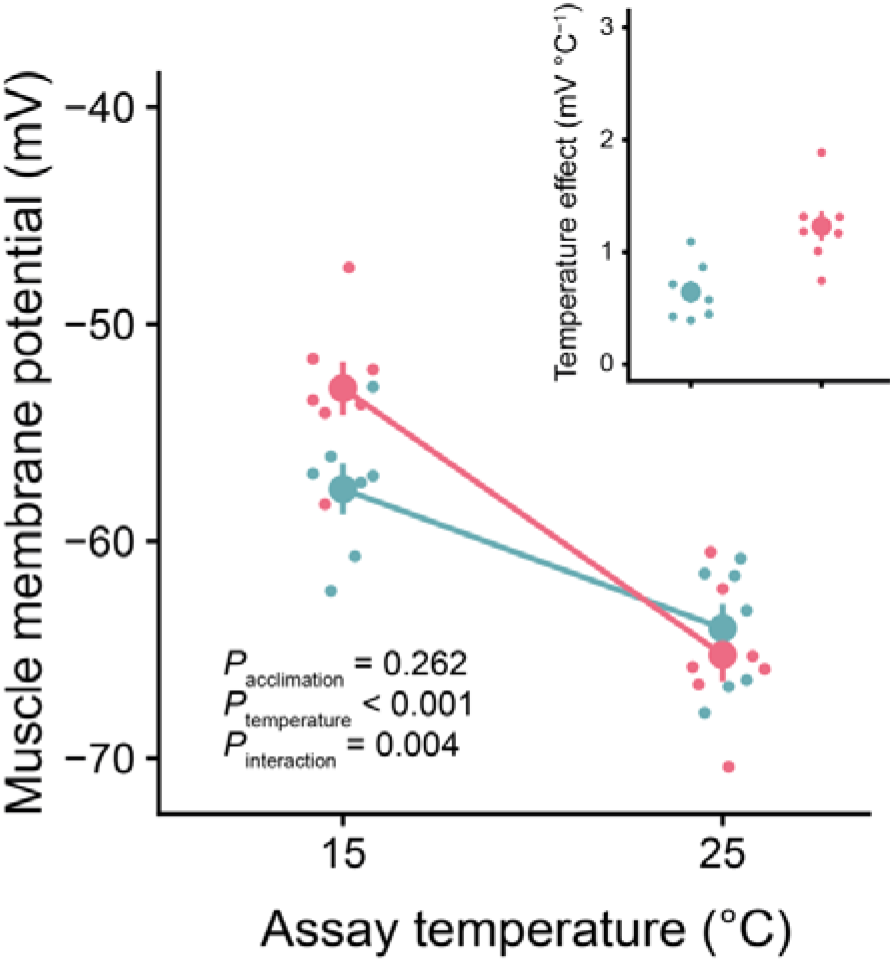
Acclimation to cold improves maintenance of the *in vivo* muscle membrane potential during cold exposure. Muscle membrane potentials were measured in the same fly at 15 and 25°C in both warm-(25°C, red) and cold-(15°C, blue) acclimated *D. melanogaster*. The insert (top-right) shows the direct effect of temperature on muscle membrane potential in both acclimation groups. Seven flies from each acclimation group were sampled in this experiment. Small circles represent individual observations.

## Discussion

### Cold acclimation alters Na^+^/K^+^-ATPase activity and thermal sensitivity in a tissue-specific manner

Given the well-documented thermal sensitivity of enzymes, Na^+^/K^+^-ATPase activity was expected to decrease with decreasing assay temperature in both the head and thorax (Hochachka and Somero, 1984). The *Q*_*10*_ values (calculated for the temperature range 15-25°C) were relatively similar to previously reported values (approximately 1.5 - 2.5 observed in mammals and crayfish; (Else et al., 1996; Zhu and Cooper, 2018). In both the head and the thorax, however, warm-acclimated flies had higher *Q*_*10*_ values (head *Q*_*10*_ = 2.15; thorax Q_10_ = 2.14) compared to those from cold-acclimated flies (head *Q*_*10*_ = 1.70; thorax *Q*_*10*_ = 1.37). These results suggest that cold acclimated flies may undergo changes in thermal sensitivity of Na^+^/K^+^-ATPase specifically in the head and thorax tissues that were not previously recorded when comparing whole animal samples (MacMillan et al., 2015c). Intriguingly, while Na^+^/K^+^-ATPase from both the thorax and head had lower thermal sensitivity, the impact of cold acclimation on the shape of the temperature-activity relationship was quite different between the two segments: While the activity of Na^+^/K^+^-ATPase from the thorax appears to have undergone a leftward shift in the curve, there instead was a wholesale reduction in Na^+^/K^+^-ATPase activity in the head (brain; Fig. 2), specifically at assay temperatures 20 and 25°C. The Na^+^/K^+^-ATPase is known to play different roles in ionoregulation in different tissues, so it is not surprising that this enzyme may respond to a temperature change in different ways in different tissues. Nonetheless, this finding should be taken into consideration, particularly because whole animal samples have been used to test for a role of this enzyme in thermal adaptation and plasticity in insects in the past (MacMillan et al., 2015c; McMullen et al., 2010; McMullen and Storey, 2008), and this approach may obscure important patterns and thereby lead to erroneous conclusions on the mechanisms underlying thermal performance. For this reason, we recommend a tissue-specific approach be used whenever possible to address similar questions in the future.

### Changes to muscle Na^+^/K^+^-ATPase activity help prevent cold-induced cell death

A decrease in thermal sensitivity with cold acclimation suggests that these tissues may be able to maintain Na^+^/K^+^-ATPase function at cooler temperatures compared to their warm acclimated conspecifics, resulting in improved cold tolerance. In the thorax (primarily muscle), maintenance of Na^+^/K^+^-ATPase activity at low temperatures would serve to maintain muscle fibre resting potential in the cold (Djamgoz and Dawson, 1988; MacMillan et al., 2014), and indeed we observed an improved ability of cold-acclimated flies to maintain muscle cell membrane potential at lower temperatures (Fig. 3), as has been observed in cold acclimated locusts (Andersen et al., 2017a) as well as cold-adapted *Drosophila* and butterfly species (Andersen et al., 2015, 2017b; Andersen and Overgaard, 2019). Given that our Na^+^/K^+^-ATPase measures maximal activity of the pump in the absence of other structures (e.g. cell membrane, cytoskeleton), our results thus support the hypothesis that thermal acclimation can prevent muscle cell depolarization, in part, through modification of the Na^+^/K^+^-ATPase itself. Depolarization of muscle cell membrane potential is no-longer thought to be directly linked to chill coma onset (Robertson et al., 2017), but activity of Na^+^/K^+^-ATPase in the muscles may be important in rebalancing K^+^ following rewarming (MacMillan et al., 2014). During a cold stress, cell depolarization directly leads to opening of voltage sensitive Ca^2+^channels that drives muscle cell death in the cold in locusts (Bayley et al., 2018), so preservation of muscle Na^+^/K^+^-ATPase activity in the cold may also directly influence chill coma recovery time as well as cell survival in the cold. Interestingly, cold acclimation was recently found to prevent cold-induced depolarization independent of changes in Na^+^/K^+^-ATPase activity or transcription (Bayley et al., 2020). Thus, while our results point towards a clear role of modulated Na^+^/K^+^-ATPase activity in maintaining muscle membrane potential during cold exposure, other mechanisms are almost certainly involved (e.g. increased muscle K^+^ permeability, see Bayley et al. 2020).

### Cold acclimation reduces temperature sensitivity of Na^+^/K^+^-ATPase in the brain but also causes lower activity at benign temperatures

In the brain, the observed decrease in thermal sensitivity of Na^+^/K^+^-ATPase would help prevent spreading depolarization (SD) in the cold (Robertson et al., 2017; Spong et al., 2016b), and may thus directly mediate the lowered chill coma onset temperature of flies acclimated to 15°C. In addition to this reduction in thermal sensitivity, we expected to see a wholesale increase in brain Na^+^/K^+^-ATPase activity in cold acclimated flies as we hypothesized this would further protect against SD in the cold. Instead, we saw the opposite effect (Fig. 2a). Thus cold-acclimated flies have a lower SD temperature despite appearing to have less, rather than more, Na^+^/K^+^-ATPase activity in their brain. With reduced Na^+^/K^+^-ATPase activity at benign temperatures, these cold acclimated flies still maintain a more positive transperineurial potential at benign temperatures. This finding suggests that reductions in Na^+^/K^+^-ATPase activity in the brain can be compensated for by other changes to epithelial physiology. The transperineurial potential is a complex trait dependent on trans- and paracellular currents, resistances and electrochemical gradients, that are maintained in part by the continuous action of ionomotive pumps (most notably the Na^+^/K^+^-ATPase). This potential is also likely to be heavily dependent on the electrochemical permeability of the glia (Treherne and Schofield, 1984). The transperineurial potential can thus be altered through changes to transcellular (e.g. channels) or paracellular (e.g. septate junction) ion permeability. While the permeability of the septate junctions that make up the blood-brain barrier has not been studied in the context of cold acclimation, the gut epithelia of *Drosophila* have longer and more convoluted septate junctions following cold acclimation (MacMillan et al., 2017), and disruption of the blood-brain barrier with 3 M urea exacerbates anoxia-induced SD in locusts (Spong et al., 2014). Therefore, adjustments to paracellular permeability in the brain could explain our results. Reduced transcellular permeability could also be driven by increased cell size, which has been observed for Malpighian tubules in cold-acclimated *D. melanogaster* (Yerushalmi et al., 2018). Lastly, a more positive transperineurial potential could be achieved *via* channel arrest mechanisms (see: Buck and Hochachka, 1993; Hochachka, 1986), or via changes to the relative ion permeabilities (specifically the K^+^ permeability, see Bayley et al., 2020). All of these mechanisms promote maintenance of cell membrane potential (and by extension the transperineurial potential) independently of the Na^+^/K^+^-ATPase and may be essential at low temperature when the pump slows down. In combination with a lower thermal sensitivity of the pump, cold-acclimated flies therefore appear able to prevent central nervous system failure (i.e. spreading depolarization onset) down to a lower temperature by 1) having superior ionoregulatory capacity in the cold and 2) relying less on the Na^+^/K^+^ ATPase for maintaining membrane polarization.

### Conclusions

Cold acclimated flies may mitigate the effects of chilling on ion homeostasis in part by decreasing the thermal sensitivity of Na^+^/K^+^-ATPase in their head (brain) and thorax (muscle). Contrary to our hypothesis, protection against SD in cold acclimated flies is likely unrelated to a wholesale increase in Na^+^/K^+^-ATPase activity, which suggests that other changes to neurophysiology may also play a role in thermal plasticity of neural function in insects. In the muscle (thorax) we observed an apparent shift in thermal optima of Na^+^/K^+^-ATPase that may be mediated by alternative transcript expression or post-translational modification of the pump and assist in chill coma recovery and muscle cell survival in the cold. A great deal of further work is required to determine the exact roles of Na^+^/K^+^-ATPase expression, localization, and function in cold acclimated and adapted insects, and future attempts should adopt a tissue-specific approach and consider the role of the membrane environment in shaping pump function under different thermal conditions.

## Supporting information

Data archive

## Data Accessibility

All data is provided as a supplementary file for review and the same file will be uploaded to a data repository (e.g. Dryad) or included as a supplementary file should the manuscript be accepted for publication.

## Author Contributions

All authors contributed to the conception and design of the study and A.C. and M.K.A. conducted the experiments. A.C., M.K.A. and H.M. analyzed the data, A.C. drafted the manuscript, and all authors edited the manuscript.

