## Supplementary materials for "Thermal acclimation alters the roles of Na^+^/K^+^-ATPase activity in a tissue-specific manner in *Drosophila melanogaster*"

**Supplementary information**

This document contains a supplementary figure for Checlock et al. (202***?***) – ” Thermal acclimation alters the roles of Na+/K+-ATPase activity in a tissue-specific manner in *Drosophila melanogaster*”

**Figure S1**

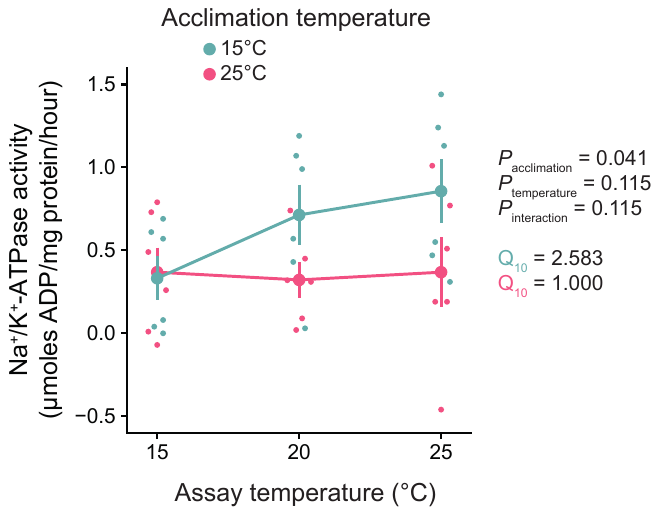

**Na^+^/K^+^-ATPase activity in the abdomen of cold- and warm-acclimated *D. melanogaster*.** Na^+^/K^+^-ATPase activity assays were performed in abdomen samples collected from 15°C (blue) and 25°C (red) acclimated flies at three temperatures (15, 20 and 25°C). Six biological replicates were sampled per acclimation and measured across the three temperatures. Small circles represent individual observations.
